# About the alignment and 3D reconstruction of sparse cryo-scanning transmission electron tomography datasets

**DOI:** 10.1101/2025.07.28.667105

**Authors:** Sylvain Trépout

**Author notes:** Corresponding author *Email address:* (Sylvain Trépout).

## Abstract

In electron microscopy, sparse imaging consists in the collection of a limited subset of the image pixels, which can be used to reduce electron beam damage. Scanning transmission electron microscopy (STEM) is particularly adapted to sparse imaging owing to the scanning nature of the method, scan patterns can be designed where fewer sample locations are targeted. However, since some of the pixels are not scanned, there is an inherent loss of information. Several algorithms were developed to reconstruct missing pixels with high fidelity. Whereas sparse imaging and missing pixel reconstruction in 2D experiments are mature methods, the application of sparse imaging in 3D scanning transmission electron tomography (STET) is rare and still under development. The main difficulty encountered in tomography studies is the tilt-series alignment, which must be accurate to ensure high-quality 3D reconstruction. Because sparse images contain only a certain portion of the original information, the images constituting sparse tilt-series might not share enough mutual information to guarantee an accurate alignment, even after missing pixel reconstruction. This work presents for the first time a thorough analysis of the fiducial alignment and reconstruction of sparse (cryo)STET tilt-series. Furthermore, the limits of sparse imaging are explored to estimate the minimum amount of information required to obtain good-quality 3D reconstructions. The use of a cryo-fixed biological sample is motivated by the fact that cryo-samples are typical highly beam-sensitive samples, and that the intricate nature and structure complexity of biological samples place them among the most difficult to reconstruct with high details.

## Introduction

In scanning transmission electron microscopy (STEM), the electron beam is focused, forming a probe at the level of the sample [1]. To create a projection image of the sample, the beam is scanned in a raster, and the electrons are detected on STEM detectors located underneath the sample. The pixels are created by integrating the number of electrons that reach the detector over a certain period of time (called the dwell time). When all pixels are scanned and the beam reaches the end of the line, the beam is rapidly positioned at the beginning of the next line to reduce damage to the sample during the beam movement. The scan of the next line does not happen subsequently after, as the beam position needs to stabilise to ensure that the correct sample coordinates are scanned. The time that occurs between the scans of two consecutive lines is called the flyback time. If the flyback is set too short, the beam does not have enough time to settle, and the left side of the image is distorted by a wavy pattern not corresponding to the actual sample structure. This scanning pattern is a historical legacy of cathode ray tube design and is used by most scan generators installed on STEM microscopes [2]. The interaction of the microscope electrons with the sample atoms is destructive and can cause damage to the sample. Electron damage results in a modification of the sample structure, preventing recovery of the genuine sample structure. Cryofixed biological samples, zeolites, and perovskites are particularly beam-sensitive samples, making their study challenging. Moreover, some imaging methods (e.g. high magnification, spectroscopy and other chemical analysis studies) are intrinsically high electron fluence methods [3]. This is the reason why development of low-fluence imaging methods is crucial for the characterisation of electron-sensitive samples.

During the 2010s, several STEM developments led to the realisation of low-fluence imaging methods. Binev *et al.* pioneered the development of computational methods, based on data compression strategies, to faithfully reconstruct missing pixels in downsampled images [4]. Then, sparse imaging was developed; the method consists in the collection of a limited subset of pixels that constitute an image, mechanically reducing electron fluence. The raster scanning pattern used in STEM is by nature compatible with sparse imaging, it just requires some pixels to be skipped during collection. Sparse imaging was pioneered in 2013 by Anderson *et al.* on a scanning electron microscope (SEM) [5]. Sparse images were collected using an external scan generator and reconstructed using a discrete cosine transform (DCT) inpainting algorithm. In 2016, Béché *et al.* were the first to use sparse imaging for electron fluence reduction purposes [6]. Sparse imaging was achieved by installing an electromagnetic shutter at the condenser level of the microscope, which was deflecting the beam towards an aperture while scanning coils conventionally scanned the beam. Sparse image reconstruction was performed using DCT inpainting. Approximately the same time, Kovarik *et al.* developed a scan generator to collect sparse images [7]. In this setup, the beam is conventionally scanned from left to right while random fluctuations of the scanning coils are imposed along the vertical axis. This solution was chosen because jumping from one sample position to another requires time for the beam to settle (quote: “time required for the beam to reach 90% of the desired position is approximately 60 *µ*s”, to compare with the dwell time of 31 *µ* used in their study), similar to when the beam flybacks during a normal raster scan. They used beta process factor analysis (BPFA), a dictionary learning algorithm, to inpaint the sparse images.

Since its inception about 10 years ago, sparse imaging is now a mature method, thanks to the development of more elaborated sparse scans and the increased efficiency of inpainting algorithms. The method is now used to characterise how electron beam damage occurs. To study the effect of electron deposition over time, smart sparse sampling schemes are designed to prevent the beam from returning to the same area too quickly, so any local effect caused by the beam has time to dissipate before the beam comes back [8, 9, 10, 11]. One of the last developments in the field is the software SenseAI [12]. By using a random walk algorithm, high sparse scanning speed can be achieved, and together with the development of an efficient GPU-based BPFA algorithm, sparse images are accurately reconstructed on-the-fly as the beam scans the sample. SenseAI offers the possibility of reducing long acquisition times by a factor of 10 or more (depending on the microscope setup). This development is invaluable for SEM, (cryo-)FIB-SEM or 4D-STEM experiments, which typically require several hours of data collection.

The works cited above refer to the study of 2D samples. Sparse imaging in 3D STEM tomography exists too, however, it is rare because it has additional challenges; sparse tomography is more complex than the cumulative sum of difficulties encountered when performing 2D sparse imaging at multiple tilt angles. The complexity comes first from the design of a tomography collection software capable of tracking an object of interest for which no complete image is collected. To my knowledge, except for the SenseAI BPFA software [12], all other reconstruction algorithms require at least several tens of seconds to reconstruct a sparse image. However, SenseAI has not been used for tomography yet. STEM tomography being already quite slow compared to its non-scanning counterpart, it is not desirable to lengthen the process even more. The second complexity, rarely mentioned and probably the most difficult one, is the correct alignment and the 3D reconstruction of sparse tilt-series.

Early sparse tomography studies used an alternative to sparse imaging that is only available in tomography, tilt-downsampling, which consists in reducing the number of tilts collected. This method can be easily and quickly implemented on any electron microscope. However, due to the limited angular information, it requires a specific 3D reconstruction algorithm, such as total variation inpainting or compressive-sensing [13]. Although tilt-downsampling proved to be advantageous for material science samples, it failed to faithfully reconstruct the intricate and complex structure of biological samples [14]. Important improvements were made when coupling tomography and image-downsampling (sparse imaging), which was found to be superior to tilt-downsampling [15, 16]. Image-downsampling outperformed tilt-downsampling in all respects, as not only the quality of the 3D reconstructions was better preserved isotropically due to the presence of the entire angular range, but higher downsampling values could also be achieved without sacrificing 3D reconstruction quality thanks to the data redundancy intrinsic to tomography. In these early sparse STEM tomography works, downsampling of the images was performed *in silico* from a conventional fully-sampled tilt-series dataset. It is only in 2019, that two tomography studies performed actual sparse sampling, the first in material sciences [17] and the second in biology [18]. For the study of the biological sample, I developed the first STEM tomography software capable of sparse image collection with automatic focusing and tracking tasks [18]. Missing pixel reconstruction and 3D reconstruction were performed using DCT inpainting [19, 20] and simultaneous iterative reconstruction technique (SIRT) [21, 22], respectively. However, in all these sparse tomography studies, whether sparse images were actually collected or generated *in silico*, the alignment quality of the sparse tilt-series was not further investigated or compared to the alignment of a conventional fully-sampled tilt-series.

The biological sample used in the 2019 sparse tomography study was a resin-embedded *Trypanosoma brucei* specimen that could sustain high electron fluence [18]. The microscope that I used at that time, a JEOL 2200FS, did not have a fast beam shutter but was equipped with a Digiscan II. This setup allowed to script the acquisition of randomly distributed pixel blocks (whose size was 16*4 pixels), as scanning single pixels was not authorised by the software (more details about the data collection in the original work). When trying to reproduce the same experiment on a cryo-sample, the sample returned systematically severely damaged by the beam after data collection. What was not visible with an electron-resilient resin sample was that the time needed for the electron beam to settle at the beginning of each collected pixel block was long (as mentioned in Kovarik *et al.* [7]) and that this strategy could not be used for an electron-sensitive cryo sample. For this reason, in 2021, I changed the scanning strategy to collect individual lines of pixels. Lines can be horizontal as in a normal raster scan or can be tilted. The collection of lines has several advantages. First of all, it does not differ much from conventional raster scanning, so its implementation is relatively easy. Then, the beam scans the sample at constant speed, so there is no need to consider any speed changes that would produce varying dwell times. Tilted lines have the potential of scanning a region of interest with a more homogeneous distribution of the electrons compared to horizontal parallel lines; however, it is more difficult to implement for the simple reason that the trajectory of the lines (angle and position) have to be pre-calculated to satisfy homogeneous distribution. Horizontal lines are trivial to implement, the only variable being the number of lines that are skipped during data collection. With colleagues, we used horizontal lines to explore the structure of an entire double-flagellated *T. brucei* almost 2 µm-thick cell, a work presented during the Microscopy and Microanalysis conference in 2021 [23] and highlighted in a newsletter journal [24]. After data collection of sparse lines, we used DCT inpainting to reconstruct the missing pixels [19, 20], Etomo to perform fiducial alignment on sparse images [25, 26], and SIRT in Tomo3D for 3D reconstruction [21, 22]. This work pioneered the use of fiducials to align sparse tilt-series. The details in the 3D volume allowed us to identify the two cell flagella, and the microtubule doublets of the axonemes were also visible inside both flagella.

These developments in tomography allow now to collect sparse STET dataset. However, the alignment and 3D reconstruction quality have still not been measured or compared to fully-sampled tilt-series. This is critical because in tomography, the alignment quality of a tilt-series determines the visibility and veracity of the details inside the 3D reconstruction. If the alignment is not precise, the reconstructed structures (e.g. membranes, filaments) will suffer strong deformation artefacts, which will affect our understanding of the sample structure. In biology, best alignments of cryo-tilt-series are achieved by tracking the positions of gold beads (GBs) used as fiducials. GBs are often used in cellular tomography as they can be easily added to the sample prior to freezing, and the wide range of commercially available GBs makes it easy to find the right GB diameter for any magnification used. However, in sparse tomography experiments, where the collected images are incomplete, the use of GBs can be challenging:

‐ Because their sizes represent just a few pixels, can GBs be accurately tracked in sparse tilt-series?
‐ How does sparse data alignment compare relative to conventional fully-sampled data alignment?
‐ And finally, does sparse data 3D reconstruction generate faithful 3D volumes?

This work presents, for the first time, a thorough analysis of the fiducial alignment of sparse cryo-STET tilt-series. This work is divided into two main parts. The first part focuses on the detection of the GBs in sparse images, to determine the sparsity limits. The second part presents a direct comparison of pairs of 3D reconstructions generated from a fully-sampled tilt-series and a sparse one, to estimate the quality of sparse 3D reconstructions.

### 1. Fiducial alignment of *in silico* sparse tilt-series

The first part of this study uses *in silico* data to investigate detection and tracking of GBs in sparse images. Sparse images were generated by removing random pixels from the images of the Etomo tutorial dataset, consisting of 64 images (1k * 1k pixels) of a *Chlamydomonas reinhardtii* cell embedded in a plastic section, on top of which, GBs were added to allow fiducial alignment [25, 26, 27]. Reconstruction of the missing pixels was performed using DCT inpainting [19, 20] as previously described [18]. First, the analysis focuses on cropped images of a single GB to investigate how accurately GBs can be detected in sparse images. Then, entire tilt-series images are analysed to study the effect of sparse data at a large scale. More information about the generation and inpainting of the sparse images can be found in the Materials and Methods section.

#### 1.1 Gold bead detection in sparse images

In this first *in silico* analysis, an image of a GB was used to generate several sparse versions of the GB. Several downsampling values were tested (from 1 to 99% of the original pixels removed) and the missing pixels were reconstructed using DCT inpainting. Figure 1 shows the distance between the detected position of the GB in the inpainted sparse images and its real position in the original image. Representative images of the downsampled GB are displayed (Figure 1, top right corner). As observed in the thumbnail images, the GB is poorly reconstructed at high downsampling (*i.e.*, over 95% of pixels removed), with significant modification of its size or shape. In some cases, the GB can disappear completely. At high downsampling values, there is a high chance that the few remaining pixels do not fall onto the GB position, leading to a poor reconstruction of the GB. As shown in the graph, when 98% of the original pixels were removed, the median distance between the reconstructed GB and its original position is ∼ 4 pixels and the 25th percentile is ∼2.5 pixels. This is an important shift for a GB of a size of 10 pixels. Between 98 and 90% downsampling, the median position of the detected GB is off by more than∼ 1 pixel and some GBs were detected more than 20 pixels away from the original position. Below 90% downsampling, the median distance is null and only a few GBs are detected at a distance of ∼1 pixel relative to the original position.

**Figure 1.**
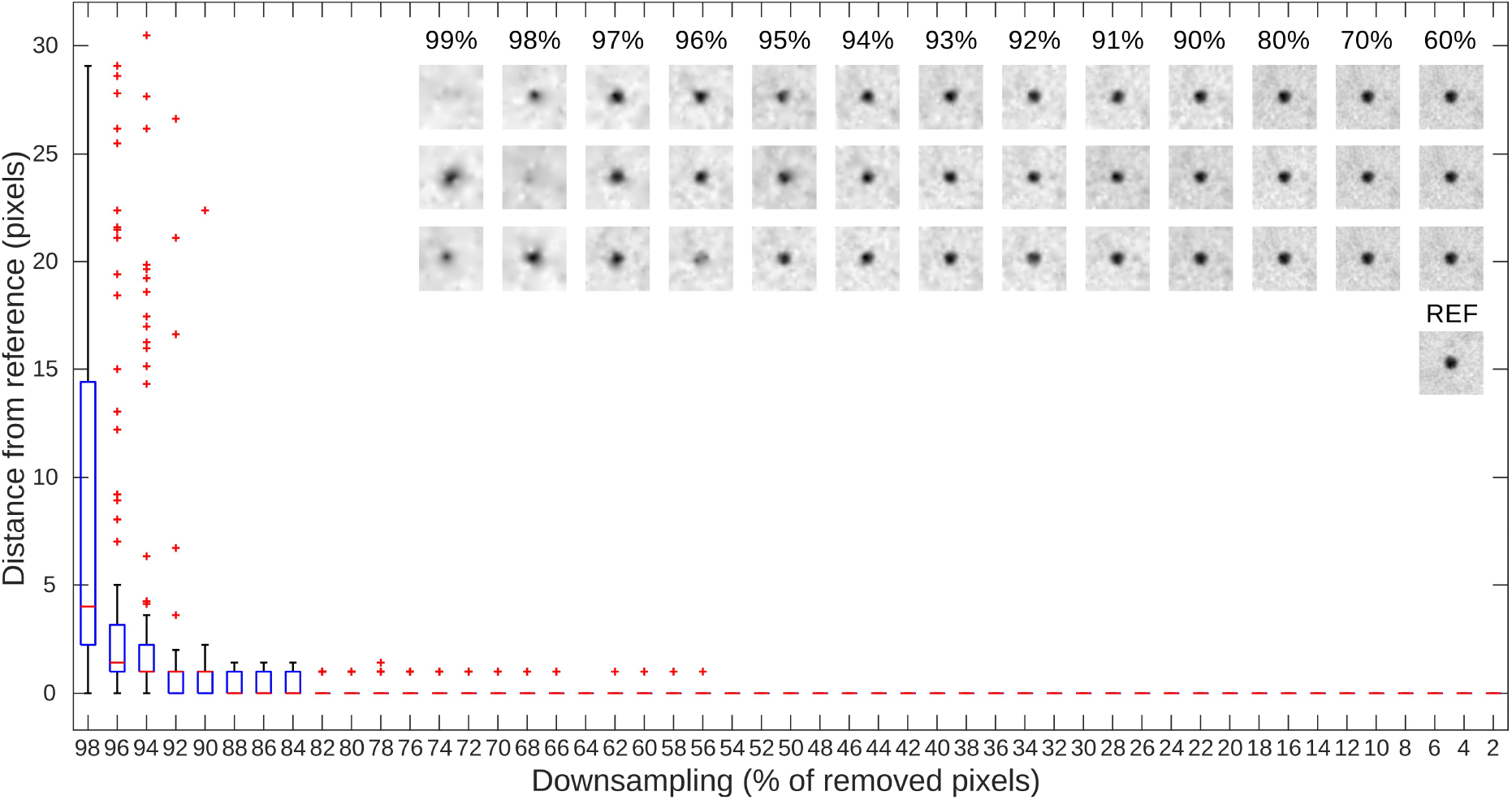
Accuracy of GB detection in inpainted sparse EM images. This plot shows the distance (in pixels) between the position of a GB detected in differently downsampled sparse images and the position of the same GB in the non-sparse reference image. Downsampling values ranged from 1 to 99% of removed pixels. To improve the visibility of the plot, only even downsampling values are represented. The median value is indicated by the horizontal red bar, the top and bottom of the blue box indicate the 75th and 25th percentiles respectively, whereas whiskers are indicated with the dashed line and the outliers are plotted with a red plus sign. The thumbnail images in the top right corner are examples of what the GBs look like after downsampling (the percentage of discarded pixels is indicated above each vertical set of 3 images) and inpainting. The reference (REF) original non-sparse image is also presented for visual comparison purposes.

These results show that when the downsampling is greater than ∼95%, the poor GB reconstruction is likely to prevent any use of a fiducial alignment strategy. Between ∼90 and ∼95% downsampling, most of the GBs are detected at the correct position, yet because some GBs are detected 20 pixels off the original position, this could translate into inaccurate fiducial alignment of the sparse tilt-series. Downsampling values lower than 90% allow a faithful reconstruction and detection of the GB, which seems suitable for fiducial alignment.

#### 1.2 Fiducial-based alignment of sparse tilt-series

To further investigate the use of a fiducial-based strategy to align entire sparse images, the downsampling/inpainting procedure was applied on the entire Etomo tutorial dataset, generating variously downsampled sparse tilt-series. For all tilt-series (*i.e.*, sparse and original ones), 20 GBs were automatically detected and tracked using Etomo [25, 26], and alignments were compared (Figure 2). Because the results on single GBs showed that the position of the GBs in sparse images could be recovered with high precision for downsampling values up to ∼90% (Figure 1), this second *in silico* analysis focuses on fewer downsampling values, aiming now at the range 90 to 99%, while including some lower values (80%, 70% and 60%) for comparison purposes. This second analysis focuses on three parameters of the fiducial-based alignment:

**Figure 2.**
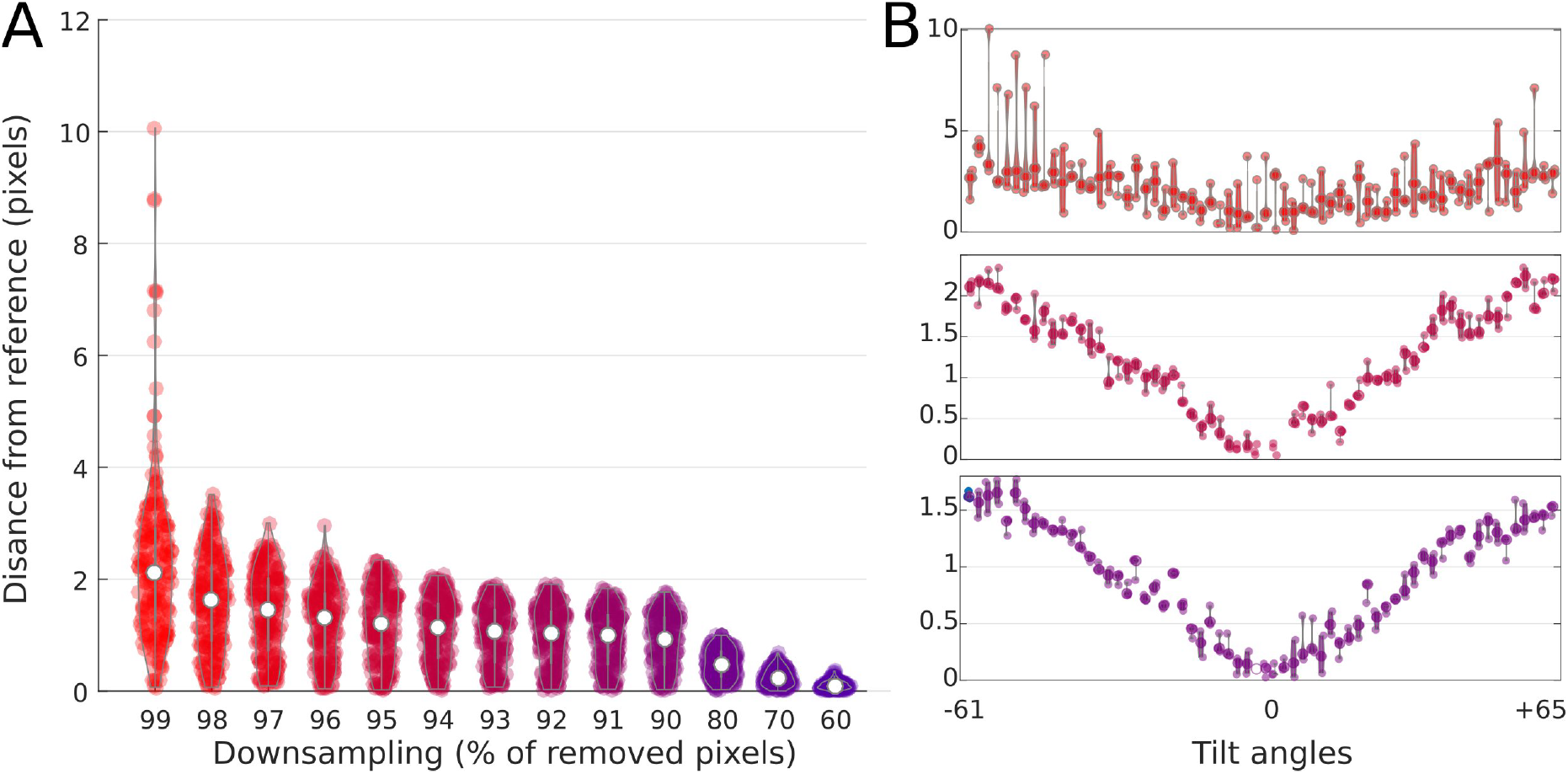
Accuracy of fiducial-based alignment of sparse tilt-series compared to the ground-truth one. The error is computed at the frame level. A) Plot showing the distances in pixels between the aligned frames of the sparse tilt-series and the original fully-sampled one. All tilt-angles are plotted together. The white dots represent the median values. The data is colour-coded to better distinguish the high downsampling values (in red) from the low ones (purple). B) Plot presenting the error in pixels as a function of the tilt angle. Only 3 downsampling values are presented (99, 95 and 90% from top to bottom). The opaque dots represent the median values.

‐ the difference between the alignment of sparse and original tilt-series.
‐ the quality of the tracking of 20 GBs.
‐ the accuracy of the GBs detection as a function of the tilt angle.

First, the analysis compares the alignment difference between the sparse and original tilt-series (Figure 2A). The difference is measured by comparing the images two by two (*i.e.*, one sparse and one original), at the same tilt angle. At 99% downsampling, the alignment error varies from 0 to 10 pixels, with a median error of ~2 pixels. The alignment error progressively decreases from 99 to 90% downsampling to reach a median error of ∼1 pixel, and the maximum error is inferior to ∼2 pixels. For lower downsampling values (80, 70 and 60%), the median error is lower than 0.5 pixel, and the maximum alignment error is inferior to 1 pixel. When represented as a function of the tilt angle, the alignment error is small for low tilt angles and important for high ones (Figure 2B). At 99% downsampling, even low-tilt images are badly aligned (Figure 2B, top panel). At 95 and 90% downsampling, high-tilt images are misaligned by ∼2 and ∼1.5 pixels, respectively, and the misalignment of low-tilt images is less than 1 pixel (Figure 2B, middle and bottom panels, respectively).

The next part of the analysis focuses on the tracking of the GBs throughout the entire tilt-series to understand why high tilt angles are more prone to bad alignment compared to low ones. Figure 3A shows the number of images on which GBs were tracked throughout the entire tilt-series, which consists of 64 frames. At 99% downsampling, the GBs are tracked throughout 49 images (median value), which includes few GBs that are tracked over ∼30 images and one GB tracked over 7 images only. At 98% downsampling, the GBs are tracked over 60 images (median value). From 96% downward, the 20 GBs are tracked over the entire dataset (64 images, median value). Plotting the number of tracked GBs per tilt-image shows that fewer GBs are successfully tracked at high tilt angles compared to low ones (Figure 3B, red plot). The important variation in the tracking of the GBs at 99% downsampling is a consequence of poor sparse reconstruction at such high downsampling. At 95% downsampling, most of the 20 GBs are tracked throughout the entire tilt range (Figure 3B, purple plot).

**Figure 3.**
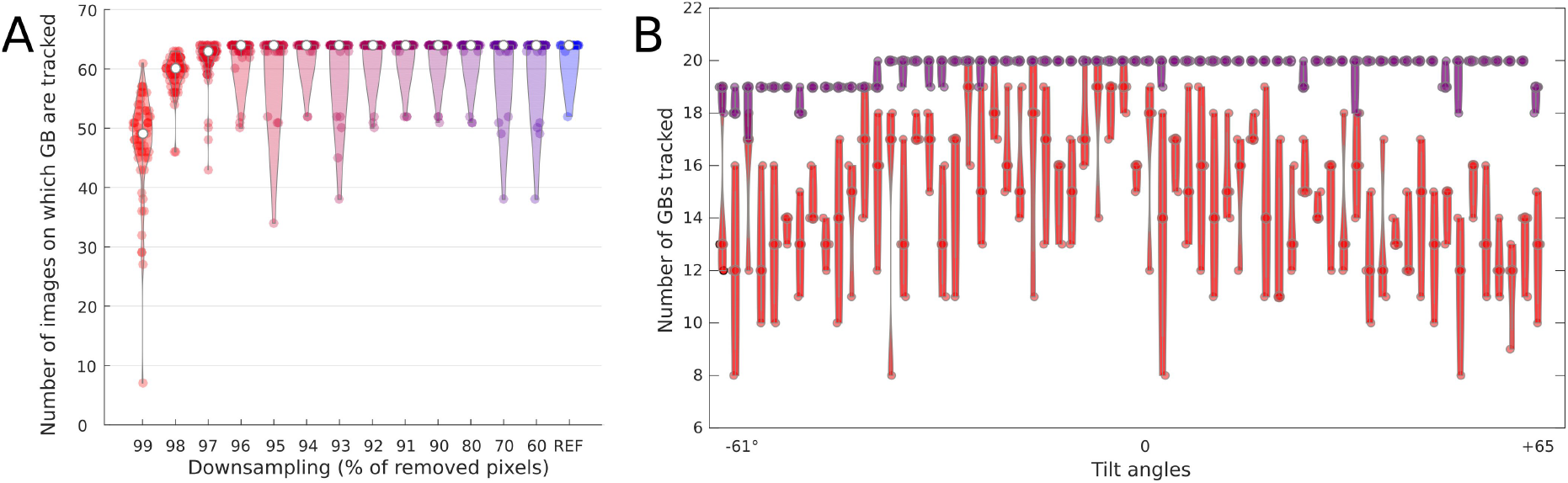
Tracking of GBs in fiducial-based alignment of sparse tilt-series. 20 GBs were automatically detected in Etomo [25, 26]. A) This plot presents the number of images on which GBs were successfully tracked throughout sparse tilt-series downsampled at various values. The tracking for the original tilt-series is also given for reference (REF). B) Plot showing the number of GBs tracked as a function of the tilt angle for 2 downsampling values (99, 95% in red and purple, respectively).

In fiducial alignment of fully-sampled data, enough GBs have to be tracked throughout the entire tiltseries to guarantee accurate alignment. However, these results show that this condition is not sufficient for sparse tilt-series, as most of the GBs are successfully tracked in the 95% downsampling experiment (Figure 3B), yet the alignment is not optimal (Figure 2B). The faithful reconstruction of sparse images is also necessary to ensure the precise picking of GBs. This appears to be more critical for high-tilt images where fewer GBs are tracked.

#### 1.3 Alignment of low-tilt and high-tilt sparse images

To further investigate the misalignment of sparse tilt-series, the X and Y components of the alignment were plotted separately for six downsampling values (99, 98, 95, 90, 80 and 60%) (Figure 4). The plots show that the X and Y components are not correlated (Figure 4, blue and red crosses, respectively). Linear regression fits show that the error along X is cumulative. Whereas the error along the X axis is important for 99% downsampling (between -4 and +5 pixels), it is less along the Y axis (between -2 and +2 pixels) (Figure 4A). For 95% downsampling, X errors are between -2 and +2 pixels, while Y errors do not exceed -0.5 or +0.5 pixels (Figure 4C). The error of the X component is still more important than the Y one at 80% downsampling (Figure 4E). There are no significant differences for the moderate downsampling value (60%) (Figure 4F).

**Figure 4.**
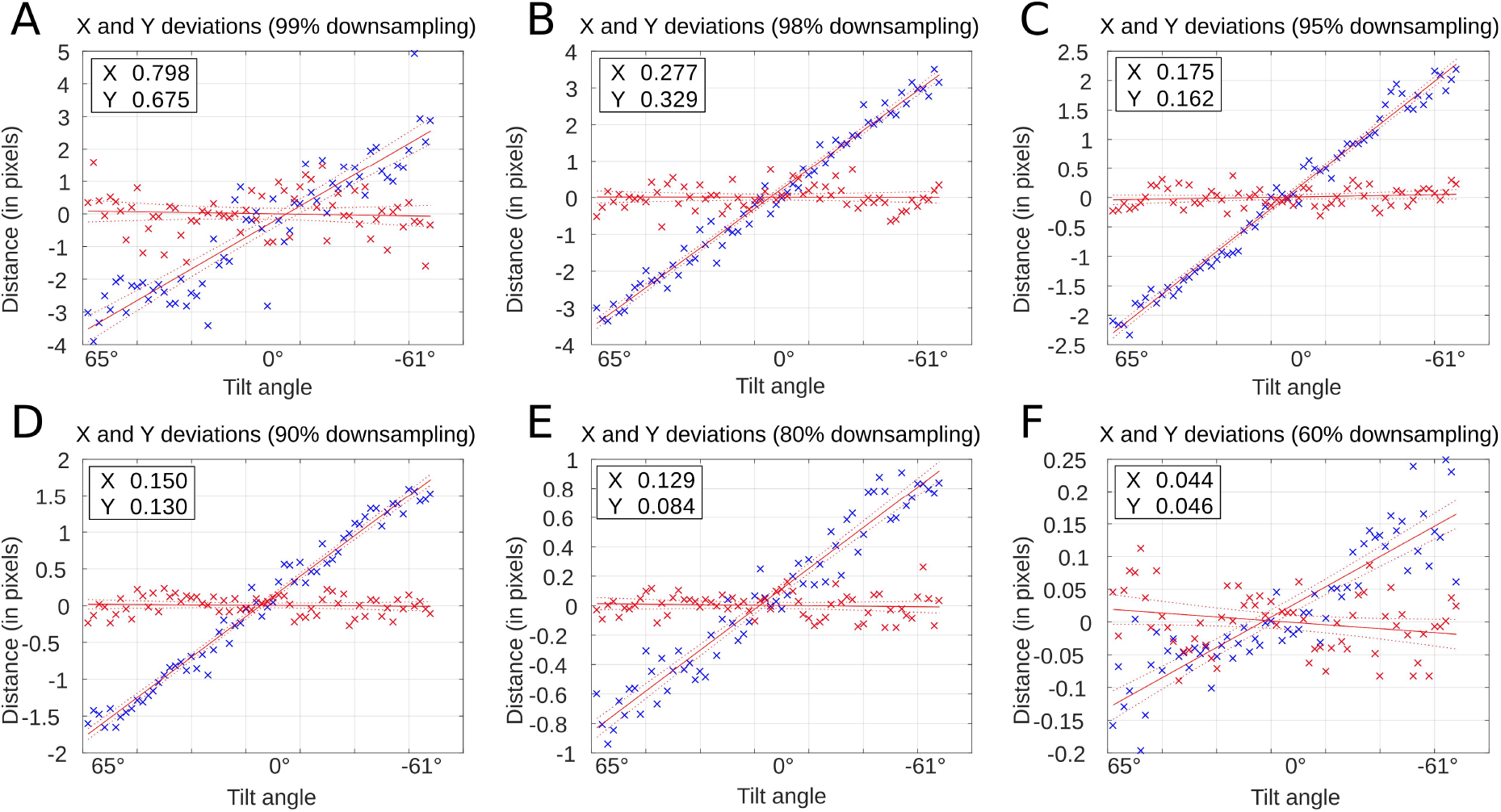
Dichotomy of X and Y shifts in fiducial-based alignment of sparse tilt-series. The plots present the X and Y discrepancies (in pixels) of differently downsampled sparse tilt-series images compared to the reference non-sparse tilt-series. A-F) The X (in blue) and Y (in red) shifts are plotted for various downsampling values (99, 98, 95, 90, 80 and 60%). Linear regression fits were computed. Values reported in the boxes represent the RMSE of the linear fits.

Note that the X (blue crosses) and Y (red crosses) errors correspond to movements parallel and perpendicular to the tilt-axis, respectively. This indicates that the misalignment of the sparse tilt-series mainly originates from the X component. Due to rotation around the tilt-axis, the shrinkage of the field of view is important along the Y axis (*i.e.*, perpendicularly to the tilt-axis). This induces a greater movement of the GBs along the Y axis compared to the X axis. Because the Y movements are much larger than the inaccuracy of the GB position determination in sparse images, the alignment along Y is driven by the large movements of the GBs perpendicularly to the tilt-axis. In contrast, the inaccuracy of the GB position determination has a much larger influence on the alignment along the X-axis. However, this hypothesis alone does not explain why the same phenomenon is not observed for low-tilt images, where the X-component error is low. In fact, the fiducial alignment software determines which GBs to pick and track in the field of view visible on the 0^°^ image. At ∼60^°^, the field of view shrinks by a factor of 2, meaning that only half of the collected pixels fall within the field of view originally visible at 0^°^. The consequence of this is that the actual downsampling value at any given tilt (DS_*α*_) is greater than the original downsampling (DS_0_) at angle 0^°^ as follows:

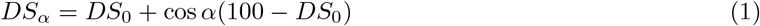

For this reason, the tracking of the fiducials at high tilts is likely to fail in experiments where the original downsampling value is already high (e.g. values higher than 80%). However, in experiments where the downsampling is modest (e.g. 60%), the tracking is expected to be accurate throughout the entire tilt-series.

#### 1.4 Discussion I

The analysis of the *in silico* data depicts some of the limits of the fiducial alignment strategy applied to sparse tilt-series. First, above 90% downsampling, the number of pixels is so low that the DCT inpainting cannot faithfully reconstruct the shape of the GBs, leading to an inaccurate detection of their positions (Figure 1) and a misalignment of the tilt-series (Figure 2). Further analysis showed that the shrinkage of the field of view at high tilts virtually increases downsampling, preventing the correct tracking of GBs, leading to misalignment of high-tilt images (Figures 3 and 4). These results have consequences on the downsampling strategy and on the data collection strategy. Downsampling values greater than ∼80% are not recommended because the alignment will not be accurate enough, especially at high tilts. Conversely, downsampling values lower than ∼80% can be used, as the alignment should be accurate. This represents a significant ∼80% reduction of the electron fluence. Furthermore, the acquisition of images at angles higher than ∼±65^°^ might not be beneficial due to their poor alignment. However, this poor alignment could be prevented by decreasing the downsampling value as the sample is tilted inside the microscope to keep the number of pixels constant in the field of view originally visible at 0^°^.

### 2. Fiducial alignment of real sparse tilt-series

The first part of this work allowed to determine the downsampling limits for fiducial alignment of sparse tilt-series generated *in silico*. In the second part of this work, real sparse tilt-series were collected on a JEOL 2200FS electron microscope at 66, 75 and 80% downsampling values, which have the potential of being accurately aligned based on the *in silico* results. Sparse data was collected using in-house scripts developed in 2019 for a previous study [18]. Instead of collecting pixel patches as in 2019, line scans were collected and combined, as first introduced in our preliminary work [23, 24]. For the sake of clarity, downsampling using randomly distributed pixels as performed in the *in silico* analysis is the best strategy as it represents the best homogeneous information distribution; however, this strategy requires a fast shutter to control the blanking of the electron beam and synchronisation with the beam scanning. A fast blanker is not a cheap and commonly installed device. Line scans can be implemented with affordable scan engines (e.g. Digiscan II, as used in this work). They are easy to handle because they are similar to raster scans except that some lines are skipped, which enables easy control of the electron deposition. After collecting sparse images using line scans, DCT inpainting was used to reconstruct the missing pixels [19, 20]. Along the sparse tilt-series, fully-sampled ones were collected and used as references in the subsequent analysis. Tilt-series were collected on *Escherichia coli* bacteria plunge-frozen on electron microscopy grids, as performed in our 2022 cryo-soft X-ray tomography study [28]. Two sparse data collection strategies were explored:

‐ low-fluence strategy: reducing the electron fluence by collecting fewer pixels and keeping the dwell time the same as for the fully-sampled tilt-series, this is an approach similar to the *in silico* analysis.
‐ high-fluence strategy: keeping the overall electron fluence the same as for the fully-sampled tilt-series by increasing the dwell time by an amount inverse of the downsampling (e.g. dwell time two times longer if two times less pixels are collected).

The low-fluence strategy is designed for the study of electron-sensitive samples, whereas the high-fluence strategy increases the signal-to-noise ratio (SNR) of the collected pixels, as could be used for methods that require an increased amount of electrons (*i.e.*, thick samples, high magnification, spectroscopy, chemical imaging methods). Electron fluences of all tilt-series are summarised in Table 1. In this second part of the work, the alignments between fully-sampled and sparse tilt-series are compared; then the analysis focuses on the visibility of bacterial structures inside the 3D reconstructions. More information about sample preparation, data collection and image analysis post-tomographic acquisition can be found in the Materials and Methods section.

**Table 1:** Data collection parameters of the fully-sampled and sparse tilt-series acquired on *E. coli*. The dwell time (expressed in µs) and the fluence (expressed in e^−^/ Å ^2^) are reported for each tilt-series. The collection parameters of the top three sparse tilt-series correspond to a low-fluence strategy. The collection parameters of the bottom three sparse tilt-series correspond to a high-fluence strategy that increases the SNR of each collected pixel.

| Tomo number | Dwell full | Fluence full | Dwell sparse | Downsampling (%) | Fluence sparse |
| --- | --- | --- | --- | --- | --- |
| Tomo110 | 4 | 60 | 4 | 66 | 20 |
| Tomo116 | 4 | 60 | 4 | 75 | 15 |
| Tomo115 | 4 | 60 | 4 | 80 | 12 |
| Tomo112 | 2 | 30 | 6 | 66 | 30 |
| Tomo113 | 2 | 30 | 8 | 75 | 30 |
| Tomo114 | 2 | 30 | 10 | 80 | 30 |

#### 2.1 Comparison of sparse and fully-sampled tilt-series alignments

After measuring the alignment quality of sparse tilt-series using *in-silico* data, the present part of this work deals with real sparse cryo-STET data. In total, six bacteria were imaged. Per bacteria, a pair of tilt-series were collected (one fully-sampled and one sparse), allowing direct comparison. Tilt-series alignment was performed using GoldDigger, a fiducial alignment algorithm that we developed recently [29]. The alignment errors as reported by the software *tiltalign* (https://bio3d.colorado.edu/imod/doc/man/tiltalign.html) are summarised in Table 2. Per bacteria, the alignment errors of the fully-sampled and the sparse tilt-series did not vary much, except for the Tomo116 which is a special case as explained below. For all the other tilt-series, the alignment difference (residual error weighted mean) is less than a pixel, as it varies from 0.1 pixel (Tomo115) up to 0.9 pixel (Tomo110). For the Tomo116, the alignment residual weighted mean errors for the fully-sampled tilt-series and the sparse one are 5.0 and 2.5 pixels, respectively. Surprisingly, the mean error is more important for the fully-sampled tilt-series. After visual inspection of the Tomo116 fully-sampled tilt-series, it appears that one of the tilt-images suffers from deformation, which was most probably caused by some microscope or environment vibration during data collection, explaining the higher alignment error.

**Table 2:**
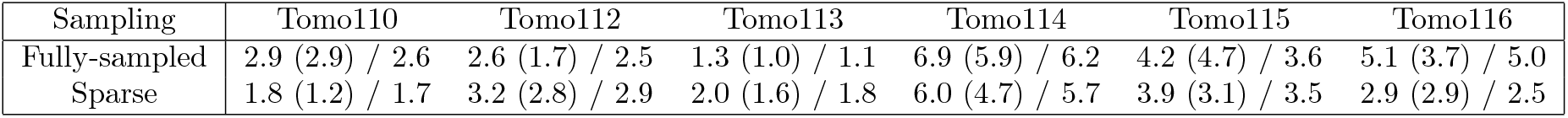
Summary of the alignment errors reported by the *tiltalign* program, part of the IMOD package [25, 26] and used in GoldDigger [29]. The reported values correspond to: residual error mean (standard deviation) / residual error weighted mean.

Based on this data, there is no evidence that fully-sampled tilt-series are better aligned than sparse ones, as the difference between the alignment errors is small (less than a pixel). Note that half of the sparse tilt-series are better aligned than the fully-sampled ones (Tomo110, Tomo114 and Tomo115). This is in agreement with what was determined in the first part of the study, the alignment quality of sparse tiltseries downsampled at 66, 75 or 80% is similar to that of fully-sampled tilt-series. It is important to note that alignment errors reported here are larger than is commonly expected for cryo-transmission electron tomography (TET). In cryo-STET, a single image is collected in several tens of seconds, as opposed to hundreds of milliseconds for movie frames in cryo-TET. During such a long collection time, the sample movement is inevitable and causes a distortion in the final image that is difficult to correct and compensate for. Methods to compensate for it exist ([30]), but are not readily applicable to cryo-STET of biological samples because they require the collection of additional images which would damage the sample. To compensate for the distortions, warping of the aligned tilt-series was computed with the software *warpalign* part of the *TomoAlign* package [31] as described in the Materials and Methods section. In fact, the alignment errors reported in Table 2 represent well-aligned cryo-STET tilt-series. In the next part of the work, the 3D reconstructions of these tilt-series are compared based on the visibility of biological structures.

### 2.2 Visibility of biological structures in sparse and fully-sampled 3D reconstructions

The visibility of biological structures is attested using 3D reconstruction images, single-image SNR analysis and plot profiles. The Materials and Methods section describes how single-image SNR values were computed. Figure 5 presents the results for the low-fluence strategy. Since the 3D reconstructions were computed using iterative methods (SIRT reconstruction in Tomo3D), the figure presents 3D volumes reconstructed at iterations 10 and 50 for both fully-sampled and sparse tilt-series, to better comprehend how the pixel values (and the visibility of biological structures) evolve as a function of reconstruction iterations. Note that in the plot profiles, vertical amber bands were added to indicate the positions of the bacteria membranes. A dent in the plot profile at this location shows that the membrane is visible in the reconstruction.

**Figure 5.**
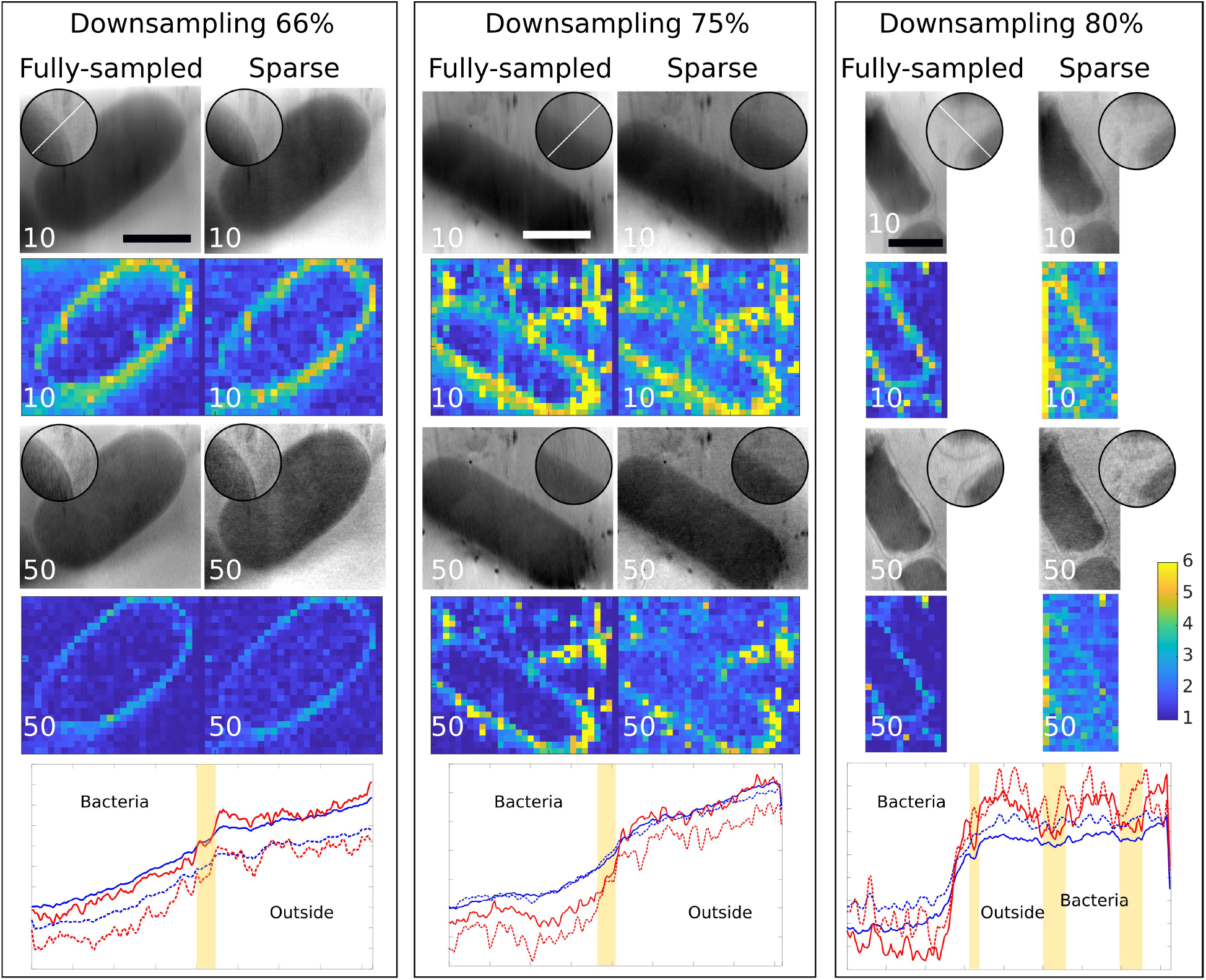
Comparison of sparse and fully-sampled 3D reconstructions in a fluence reduction strategy. Three dose reduction strategies were tested, 66, 75 and 80% downsampling. Z-slices of the different 3D reconstructions of bacteria are presented. First row, 10 SIRT iterations. Second row, single-image SNR analysis of first row. Third row, 50 SIRT iterations. Fourth row, single-image SNR analysis of third row. Round inserts are blown-up images of the reconstructions, with white lines indicating where the plot profiles were measured. Bottom row, plot profiles (blue is 10 SIRT iterations, red is 50 SIRT iterations, and dotted represent sparse data). The amber vertical bands represent the positions of the membranes, separating the inside of the bacteria from the outside. Scale bars are 1 *µ*m. The colour-coded scale of the SNR values is indicated on the right side of the figure.

As a general observation, all reconstructions computed with 10 iterations only are blurry, typical of reconstructions that have not converged yet (Figure 5, top rows). In the fully-sampled data, the contours of the bacteria in particular have a strong SNR. For the sparse data, the contours of the bacteria also have a strong SNR, but the SNR of the background (inside and outside the bacteria) increases as the downsampling gets higher (Figure 5, second rows). After 50 iterations, all fully-sampled reconstructions are crisper, and membranes become visible and sharp. For sparse data after 50 iterations, the reconstructions quality improved compared to 10 iterations, but the image texture gets grainier as the downsampling value increases (Figure 5, third rows). The contours of the fully-sampled bacteria still have a strong SNR, though lower than at 10 iterations, but the contours are thinner and the background SNR is homogeneous and low. For the sparse data, the SNRs of the bacteria contours also become thinner, but the background SNR remains high, especially at 80% downsampling (Figure 5, fourth rows). Based on the plot profiles, the bacteria membranes are barely visible for all reconstructions computed with 10 iterations only (Figure 5, bottom rows), in agreement with the reconstruction images (Figure 5, top rows). At 66% downsampling (red dotted line), there is little difference with the fully-sampled data (red plain line). At 75% downsampling, the plot profile starts becoming noisy (presence of numerous dents and peaks in the dotted red plot profile) which are absent in the plot profile of fully-sampled data (plain red plot) and are not representative of any structure in the reconstruction. This noise is reminiscent of the grainy texture visible in the images. At 80% downsampling, the noise becomes even more important, and the dents no longer match the membrane locations indicated by the amber bands.

Fully-sampled data are reference data. The observations for reference data are as follows:

‐ blurry images and high SNR around object borders (e.g. membranes and GBs) in early SIRT iterations.
‐ images better defined and low SNR for areas depleted of object borders in late SIRT iterations.

The observations on sparse data are different, in particular at high downsampling values. At 66% downsampling, the reconstruction images, the distribution of the SNR values and the densities in the plot profiles are almost identical to those of the fully-sampled data. However, when downsampling increases (75 and 80%), a grainy texture appears in the images, which induces a higher SNR in areas exempt from any structures, and the plot profiles become so noisy that the membrane positions are no longer visible.

Figure 6 presents the results for the high-fluence strategy. For the fully-sampled reference data, the results are strictly identical, confirming the observations made in the low-fluence strategy section. For sparse data, the reconstructions with 10 iterations are blurry (Figure 6, top rows), and more iterations are necessary to increase the visibility of the structures (Figure 6, third rows). The grainy texture of the images is still present in the sparse reconstructions, however, this does not hamper the visibility of the membranes. The evolution and distribution of the SNR values are identical to what was observed for the low-fluence strategy (Figure 6, second and fourth rows). The visibility of the membranes in the plot profiles is confirmed by the dents located inside the amber bands (Figure 6, bottom rows).

**Figure 6.**
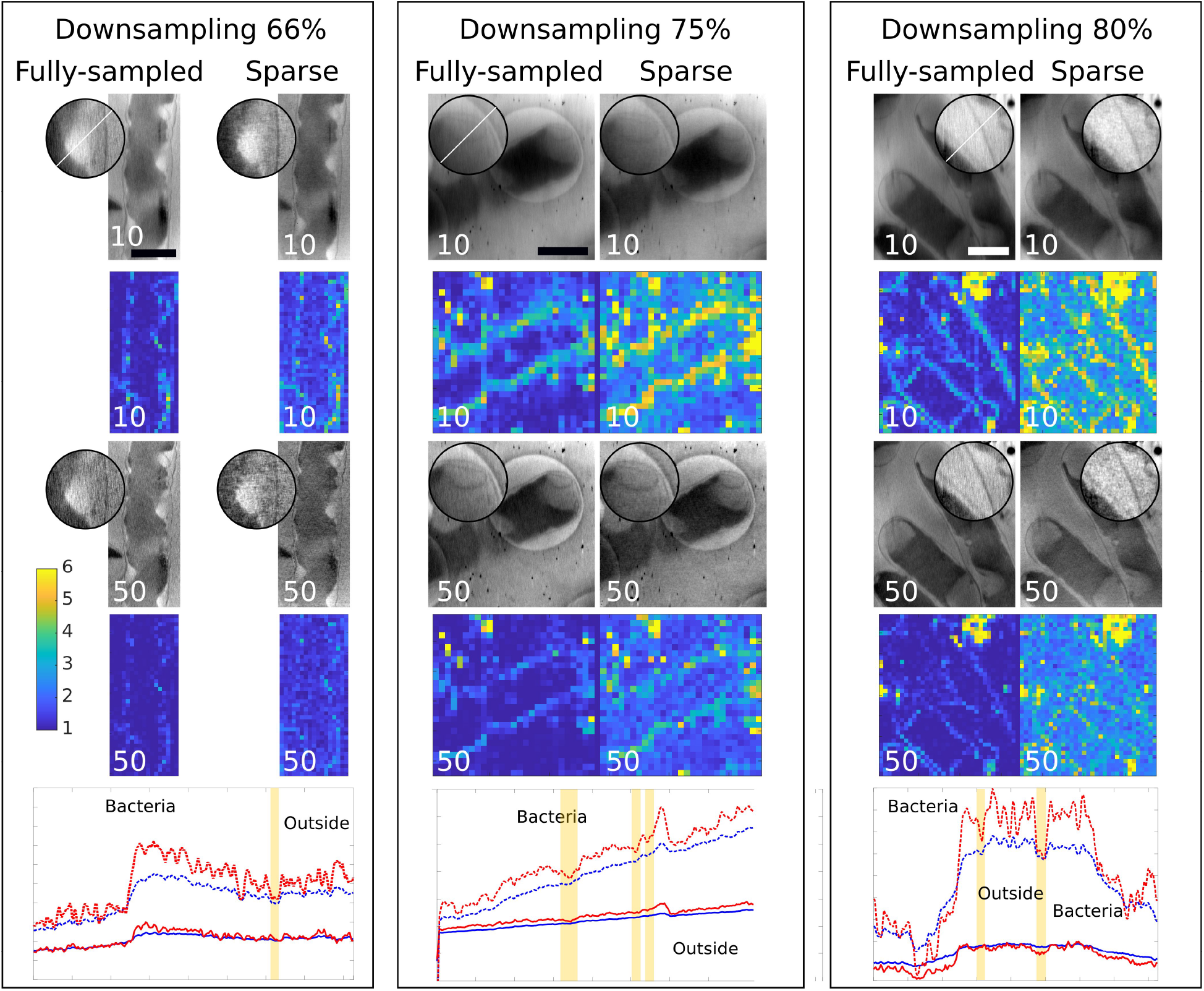
Comparison of sparse and fully-sampled 3D reconstructions in a high fluence strategy. Three downsampling were tested, 66, 75 and 80%. Z-slices of the different 3D reconstructions of bacteria are presented. First row, 10 SIRT iterations. Second row, single-image SNR analysis of first row. Third row, 50 SIRT iterations. Fourth row, single-image SNR analysis of third row. Round inserts are blown-up images of the reconstructions, with white lines indicating where the plot profiles were measured. Bottom row, plot profiles (blue is 10 SIRT iterations, red is 50 SIRT iterations, and dotted represent sparse data). The amber vertical bands represent the positions of the membranes, separating the inside of the bacteria from the outside. Scale bars are 1 *µ*m. The colour-coded scale of the SNR values is indicated on the left side of the figure.

### 2.3 Discussion II

In the second part of this work, actual sparse tilt-series were acquired, using data collection parameters determined based on the results of the *in silico* analysis. To ensure accurate alignment of the tilt-series, the highest downsampling value was 80%. The comparison of alignments confirmed that at such downsampling values, no difference could be measured between sparse tilt-series and fully-sampled ones (Table 2). Interestingly, this shows that the results measured in the *in silico* data can be directly transposed to the real data, suggesting that the beam position during the scanning of sparse lines is remarkably stable. Indeed, a criticism of using *in silico* sparse data is that it does not account for potential beam defects during scanning in unconventional ways. The analysis of 3D reconstructions showed that the low-fluence 75 and 80% downsampled datasets possessed a visibly important grainy texture which hampered the visibility of the bacteria membranes, in the images and in the plot profiles. Only the almost artefact-free 66% downsampling strategy seems appropriate for structural studies. With the high-fluence strategy, the result is totally different. Although reconstruction artefacts are still visible, the visibility of bacteria membranes is maintained in the 3D volumes for all downsampling values. Moreover, the presence of the membranes in the plot profiles is better defined than in the fully-sampled data. The high-fluence strategy as it was designed consists in extending the dwell time. This results in collected pixels with increased SNR, which appears to be beneficial for the quality of the 3D reconstructions.

## 3. Conclusion

This work presents for the first time a thorough analysis of the fiducial alignment and reconstruction of sparse (cryo-)STET tilt-series. Downsampling values up to 80% guarantee accurate alignment of the tiltseries, as tested on *in silico* sparse data. This is less than downsampling values classically used in 2D studies, but the difference is explained by the fact that tilting the sample inside the microscope shrinks the field of view, virtually increasing the downsampling up to ∼90% (number of collected pixels reduced by a factor of 2 at*±*60^°^). This significantly decreases the electron fluence required to image beam-sensitive samples, paving the way for new structural studies in both material and biological sciences. Downsampling values greater than ∼70% seem to generate a patchy grainy pattern inside the 3D reconstructions, which would have to be investigated further, as this pattern could result from the use of a regular sparse sampling (scan lines were not randomly distributed). The use of other inpainting algorithms, in particular BPFA, could meliorate the inpainting quality, improving the overall alignment and 3D reconstruction. Using sparse scanning to increase the dwell time per collected pixel produced highly contrasted 3D reconstructions, confirming that this strategy previously used in 2D spectroscopy [32] is extendable to other 3D high-fluence methods. As 2D sparse scanning software are now freely or commercially available, extension to tomography is the next step. Recent efforts to implement STEM developments in SerialEM [33, 34, 35] demonstrate how new methods can be rapidly and widely made available thanks to community-supported resources.

## 4. Materials and methods

### 4.1 Generation of in silico sparse images and tilt-series

*In silico* sparse images and tilt-series were generated from the Imod tutorial dataset (https://bio3d.colorado.edu/imod/doc/etomoTutorial.html). The dataset images were first downsampled (removal of a certain number of pixels at random locations), and then inpainted using the DCT algorithm developed by Garcia [19, 20]. All computing steps were performed in MATLAB [36].

### 4.2. Gold bead detection

In the first *in silico* experiment (Figure 1), an image of a GB was extracted from the Imod tutorial dataset. Different downsampling were performed, removing between 1 and 99% of the original pixels. Since the positions of the removed pixels were picked randomly during downsampling, each downsampling was repeated 100 times to generate 100 different datasets, producing statistical significance. After DCT inpainting, the program *imodfindbeads* (https://bio3d.colorado.edu/imod/doc/man/imodfindbeads.html) was used to find the centre of the GB. Figure 1 was generated in MATLAB [36], and shows the distance in pixels between the detected centre of the sparse GB and the centre GB in the original reference image.

### 4.3 Comparative alignment of tilt-series

In the second *in silico* experiment (Figure 2), the entire field of view of the Imod tutorial dataset was used. At each downsampling value, three different random pixel removals were computed and subsequently followed by DCT inpainting, generating three independent tilt-series per downsampling value for statistical analysis of the results. Tilt-series alignment was performed in Etomo, automatically tracking 20 GBs, and no manual correction of the fiducial locations was performed. The alignment of each sparse tilt-series was compared frame-by-frame to the reference original tilt-series. This was achieved by extracting and comparing the X and Y shift values present in the Etomo .xf files. Figure 2 was generated in MATLAB [36], and presents the distance in pixels between the aligned sparse frames and the aligned reference frames. In Figure 4, the X and Y shift values are plotted separately. The linear regression models were fitted to the data using the *fitlm* function in MATLAB [36]. The dotted lines encompass the 95% confidence interval. Values reported in the boxes represent the RMSE of the linear fit.

### 4.4 Analysis of gold bead tracking

In the third *in silico* experiment (Figure 3), the tracking of the 20 GBs and their positions were analysed. This was achieved by extracting the information contained in the Etomo .fid file using the command *imodinfo* (https://bio3d.colorado.edu/imod/doc/man/imodinfo.html). Figure 3 was generated in MATLAB [36], and presents the number of images on which GBs are tracked, and the number of GBs tracked per tilt angle.

### 4.5 Sample preparation

The sample used in this work is the same as that studied in a previous work [28]. Briefly, the bacterial strain used is *E. coli* MG1655. Bacteria were grown in LB medium until they reached an OD_600_ of 1.5. Prior to deposition on the EM grids, 10 µL of bacteria culture were mixed with 5 µL of 20 nm GBs used as fiducial markers and 5 µL of M9 media. A 5 µL drop of the bacteria and GB mix was deposited on carbon-coated gold R2/1 Quantifoil grids (Quantifoil Micro Tools, Grosslobichau, Germany), previously glow-discharged using a PELCO EasiGlow (Ted Pella, Inc., Redding, CA, USA). Grids were manually blotted and plunge-frozen in liquid ethane cooled at -174°C by liquid nitrogen using a Leica EM-CPC (Leica Microsystems, Wetzlar, Germany).

### 4.6 Cryo-STET data collection, alignment and reconstruction

Cryo-grids were mounted on a Gatan 914 high-tilt cryo-holder (Gatan, Pleasanton, CA, USA) and loaded into a JEOL 2200FS 200kV FEG electron microscope (JEOL, Tokyo, Japan). Cryo-STET datasets were collected in bright-field mode using a 20 µm condenser aperture at 40 cm camera length. The different dwell times used in this work are summarised in Table 1. The magnification used was 40,000x (corresponding pixel size is 2 nm). Cryo-STET and sparse cryo-STET datasets (image size 3000 * 3000 pixels) were collected using scripting in Digital Micrograph, as in previous experiments [18]. Tilt-series were collected between -65^°^ and +65^°^ using 3^°^ tilt increments. The total electron fluences per tilt-series are summarised in Table 1. Sparse tilt-series were collected as line scans, individually stored as they were acquired. After assembling the individual line scans to reconstitute the image, DCT inpainting was performed using 250 iterations in MATLAB [36]. Tilt-series alignment was performed using GoldDigger [29]. GoldDigger employs several Imod programs to iteratively detect fiducials and align the tilt-series. Picking of the GBs was manually verified to achieve the best alignment possible for all tilt-series. This task was more difficult for the highly downsampled sparse tilt-series as some GBs initially tracked in the low-tilt images were no longer visible in high-tilt images. Since data collection in STEM is slow (e.g. few tens of seconds per image), there is a significant sample motion during collection of an entire tilt-series. To compensate for the sample motion, tilt-series alignment was performed by warping the images using the program *tomowarpalign* [37, 38, 31]. Warping process requires a file containing the coordinates of GBs throughout the entire tilt-series. GoldDigger generates such a .fid file, using the Imod program *tiltalign* (https://bio3d.colorado.edu/imod/doc/man/tiltalign.html). 3D SIRT reconstructions were computed with Tomo3D [21, 22] using the aligned warped tilt-series as input.

### 4.7 Extracting residual and mean alignment errors

Table 2 contains residual and mean alignment errors extracted from the align.log files, generated by GoldDigger using the Imod program *tiltalign*.

### 4.8 Image analysis

3D reconstruction images (80 nm thick) and plot profiles presented in Figures 5 and 6 were generated in ImageJ [39]. SNR values were calculated as described in Algorithm 1, based on a single-image SNR method developed by Thong *et al.* [40]. Briefly, using this method, the SNR of an image can be calculated after recovering the value of a fitted virtual point, positioned at the centre of the auto-correlation function. The SNR values were calculated on small cubes of 25^3^ pixels, inside the entire 3D reconstructions. Images showing SNR values represent the centre slices of the 3D reconstructions and were generated in MATLAB [36]. All figures were generated in Inkscape (https://inkscape.org/).

#### Algorithm 1

Single-image SNR

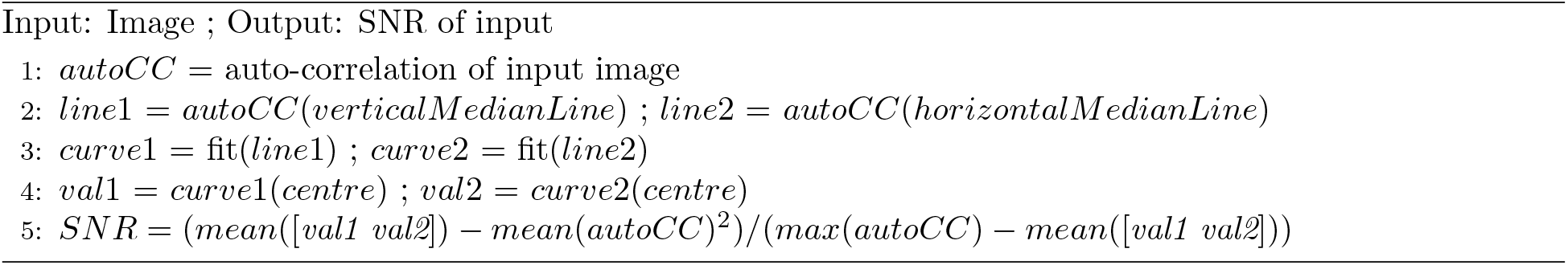

### 4.9 Data availability

All fully-sampled and sparse 3D SIRT reconstructions presented in Figures 5 and 6 were deposited in the Electron Microscopy DataBase (EMDB accession number EMD-71629).

## Acknowledgments

This work was funded by the French Agence Nationale de la Recherche, project C-STET-4-E-Cells (S.T., grant ANR-21-CE42-0001). Data were collected on the JEOL 2200FS at Institut Curie of Orsay, France, an equipment part of the PICT-IBiSA.

